# A unified analytic framework for prioritization of non-coding variants of uncertain significance in heritable breast and ovarian cancer

**DOI:** 10.1101/031419

**Authors:** Eliseos J. Mucaki, Natasha G. Caminsky, Ami M. Perri, Ruipeng Lu, Alain Laederach, Matthew Halvorsen, Joan HM. Knoll, Peter K. Rogan

## Abstract

**Background:** Sequencing of both healthy and disease singletons yields many novel and low frequency variants of uncertain significance (VUS). Complete gene and genome sequencing by next generation sequencing (NGS) significantly increases the number of VUS detected. While prior studies have emphasized protein coding variants, non-coding sequence variants have also been proven to significantly contribute to high penetrance disorders, such as hereditary breast and ovarian cancer (HBOC). We present a strategy for analyzing different functional classes of non-coding variants based on information theory (IT).

**Methods:** We captured and enriched for coding and non-coding variants in genes known to harbor mutations that increase HBOC risk. Custom oligonucleotide baits spanning the complete coding, non-coding, and intergenic regions 10 kb up- and downstream of *ATM, BRCA1, BRCA2, CDH1, CHEK2, PALB2,* and *TP53* were synthesized for solution hybridization enrichment. Unique and divergent repetitive sequences were sequenced in 102 high-risk patients without identified mutations in *BRCA1/2.* Aside from protein coding changes, IT-based sequence analysis was used to identify and prioritize pathogenic non-coding variants that occurred within sequence elements predicted to be recognized by proteins or protein complexes involved in mRNA splicing, transcription, and untranslated region (UTR) binding and structure. This approach was supplemented by *in silico* and laboratory analysis of UTR structure.

**Results:** 15,311 unique variants were identified, of which 245 occurred in coding regions. With the unified IT-framework, 132 variants were identified and 87 functionally significant VUS were further prioritized. We also identified 4 stop-gain variants and 3 reading-frame altering exonic insertions/deletions (indels).

**Conclusions:** We have presented a strategy for complete gene sequence analysis followed by a unified framework for interpreting non-coding variants that may affect gene expression. This approach distills large numbers of variants detected by NGS to a limited set of variants prioritized as potential deleterious changes.

## BACKGROUND

Advances in NGS have enabled panels of genes, whole exomes, and even whole genomes to be sequenced for multiple individuals in parallel. These platforms have become so cost-effective and accurate that they are beginning to be adopted in clinical settings, as evidenced by recent FDA approvals [1, 2]. However, the overwhelming number of gene variants revealed in each individual has challenged interpretation of clinically significant genetic variation [3–5].

After common variants, which are rarely pathogenic, are eliminated, the number of VUS in the residual set remains substantial. Assessment of pathogenicity is not trivial, considering that nearly half of the unique variants are novel, and cannot be resolved using published literature and variant databases [6]. Furthermore, loss-of-function variants (those resulting in protein truncation are most likely to be deleterious) represent a very small proportion of identified variants. The remaining variants are missense and synonymous variants in the exon, single nucleotide changes, or in frame insertions or deletions in intervening and intergenic regions. Functional analysis of large numbers of these variants often cannot be performed, due to lack of relevant tissues, and the cost, time, and labor required for each variant. Another problem is that *in silico* protein coding prediction tools exhibit inconsistent accuracy and are thus problematic for clinical risk evaluation [7–9]. Consequently, 90% of HBOC patients receiving genetic susceptibility testing will receive an inconclusive or uncertain result [10].

One strategy to improve variant interpretation in patients is to reduce the full set of variants to a manageable list of potentially pathogenic variants. Evidence for pathogenicity of VUS in genetic disease is often limited to amino acid coding changes [11, 12], and mutations affecting splicing, transcription activation, and mRNA stability tend to be underreported [13–19]. Splicing errors are estimated to represent 15% of disease-causing mutations [20], but may be much higher [21, 22]. The impact of a single nucleotide change in a recognition sequence can range from insignificant to complete abolition of a protein binding site. The complexity of interpretation of non-coding sequence variants benefits from computational approaches [23] and direct functional analyses [24–28] that may each support evidence of pathogenicity.

*Ex vivo* transfection assays developed to determine the pathogenicity of VUS predicted to lead to splicing aberrations (using *in silico* tools) have been successful in identifying pathogenic sequence variants [29, 30]. IT-based analysis of splicing variants has proven to be robust and accurate at analyzing splice site (SS) variants, including splicing regulatory factor binding sites (SRFBSs), and in distinguishing them from polymorphisms in both rare and common diseases [31]. However, IT can be applied to any sequence recognized and bound by another factor [32], such as with transcription factor binding sites (TFBSs) and RNA-binding protein binding sites (RBBSs). IT is used as a measure of sequence conservation and is more accurate than consensus sequences [33]. The individual information (*R_i_*) of a base is related to thermodynamic entropy, and therefore free energy of binding, and is measured on a logarithmic scale (in bits). By comparing the change in information (Δ*R_i_*) for a nucleotide variation of a bound sequence, the resulting change in binding affinity is ≥ 2^Δ^*^Ri^*, such that a 1 bit change in information will result in at least a 2-fold change in binding affinity [34].

IT measures nucleotide sequence conservation and does not provide information on effects of variants on mRNA secondary (2°) structure, nor can it accurately predict effects of amino acid sequence changes. Other *in silico* methods have attempted to address these deficiencies. For example, Halvorsen et al. (2010) introduced an algorithm called SNPfold, which computes the potential effect of a single nucleotide variant (SNV) on mRNA 2° structure [15]. Predictions made by SNPfold can be tested by the SHAPE assay (Selective 2’-Hydroxyl Acylation analyzed by Primer Extension) [35], which provides evidence for sequence variants that lead to structural changes in mRNA by detection of covalent adducts in mRNA.

The ramifications for better interpretation of VUS are particularly relevant for HBOC [36]. Although linkage studies suggest approximately 85% of high-risk families have deleterious variants in *BRCA1* and *BRCA2,* less than half have identified pathogenic mutations [37]. This implies that deleterious variants lie in untested regions of *BRCA1/2,* untested genes, or are unrecognized [38, 39]. Consequently, VUS in *BRCA1/2* greatly outnumber known deleterious mutations [40].

Here, we develop and evaluate IT-based models to predict potential non-coding sequence mutations in SSs, TFBSs, and RBBSs in 7 genes sequenced in their entirety in 102 HBOC patients who did not exhibit known *BRCA1/2* coding mutations at the time of initial testing. The genes are: *ATM, BRCA1, BRCA2, CDH1, CHEK2, PALB2,* and *TP53,* and have been reported to harbor mutations that increase HBOC risk [41–63]. We apply these IT-based methods to analyze variants in the complete sequences of coding, non-coding, and up- and downstream regions of the 7 genes. In this study, we established and applied a unified IT-based framework, first filtering out common variants, then to “flag” potentially deleterious ones. Then, using context-specific criteria and information from the published literature, we prioritized likely candidates.

## METHODS

### Design of Tiled Capture Array for HBOC Gene Panel

Nucleic acid hybridization capture reagents designed from genomic sequences generally avoid repetitive sequence content to avoid cross hybridization [64]. Complete gene sequences harbor numerous repetitive sequences, and an excess of denatured C_0_t-1 DNA is usually added to hybridization to prevent inclusion of these sequences [65]. RepeatMasker software completely masks all repetitive and low-complexity sequences [66]. We increased sequence coverage in complete genes with capture probes by enriching for both single-copy and divergent repeat (> 30% divergence) regions, such that, under the correct hybridization and wash conditions, all probes hybridize only to their correct genomic locations [64]. This step was incorporated into a modified version of Gnirke and colleagues’ (2009) in-solution hybridization enrichment protocol, in which the majority of library preparation, pull-down, and wash steps were automated using a BioMek^®^ FXP Automation Workstation (Beckman Coulter, Mississauga, Canada) [67].

Genes *ATM* (RefSeq: NM_000051.3, NP_000042.3), *BRCA1* (RefSeq: NM_007294.3, NP_009225.1), *BRCA2* (RefSeq: NM_000059.3, NP_000050.2), *CDH1* (RefSeq: NM_004360.3, NP_004351.1), *CHEK2* (RefSeq: NM_145862.2, NP_665861.1), *PALB2* (RefSeq: NM_024675.3, NP_078951.2), and *TP53* (RefSeq: NM_000546.5, NP_000537.3) were selected for capture probe design by targeting single copy or highly divergent repeat regions (spanning 10 kb up- and downstream of each gene relative to the most upstream first exon and most downstream final exon in RefSeq) using an *ab initio* approach [64]. If a region was excluded by *ab initio* but lacked a conserved repeat element (i.e. divergence > 30%) [66], the region was added back into the probe-design sequence file. Probe sequences were selected using PICKY 2.2 software [68]. These probes were used in solution hybridization to capture our target sequences, followed by NGS on an Illumina Genome Analyzer IIx (**Supplementary Methods - Additional File 1**).

Genomic sequences from both strands were captured using overlapping oligonucleotide sequence designs covering 342,075 nt among the 7 genes (**Figure 1**). In total, 11,841 oligonucleotides were synthesized from the transcribed strand consisting of the complete, single copy coding, and flanking regions of *ATM* (3,513), *BRCA1* (1,587), *BRCA2* (2,386), *CDH1* (1,867), *CHEK2* (889), *PALB2* (811), and *TP53* (788). Additionally, 11,828 antisense strand oligos were synthesized (3,497 *ATM,* 1,591 *BRCA1,* 2,395 *BRCA2,* 1,860 *CDH1,* 883 *CHEK2,* 826 *PALB2,* and 776 *TP53*).

**Figure 1.**
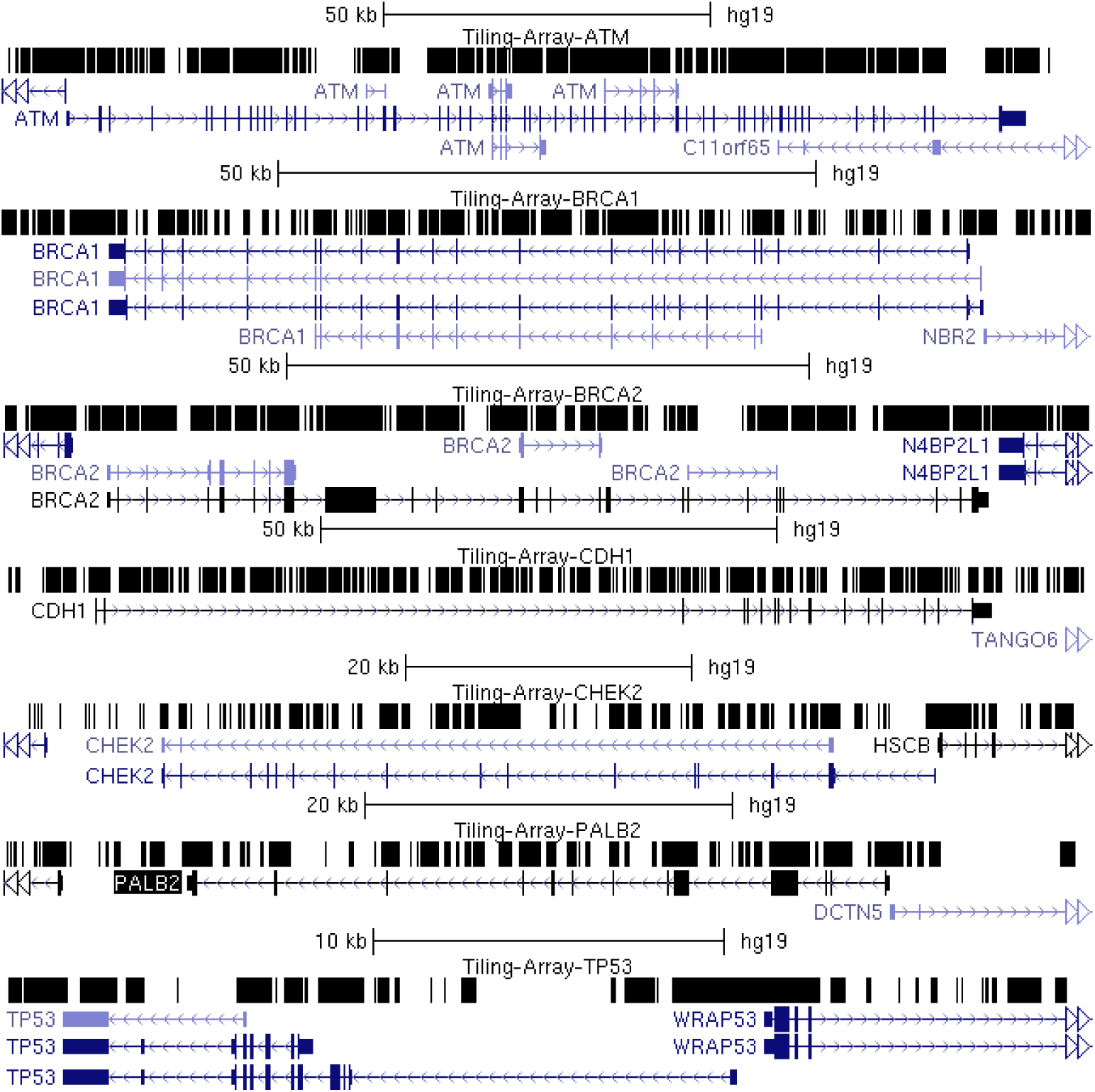
Capture Probe Coverage over Sequenced Genes. The genomic structure of the 7 genes chosen are displayed with the UCSC Genome Browser. Top row for each gene is a custom track with the “dense” visualization modality selected with black regions indicating the intervals covered by oligonucleotide capture reagent. Regions without probe coverage contain conserved repetitive sequences or correspond to paralogous sequences that are unsuitable for probe design.

For regions lacking probe coverage (of ≥ 10 nt, N=141; 8 in *ATM,* 26 in *BRCA1,* 10 in *BRCA2,* 29 in *CDH1,* 36 in *CHEK2,* 15 in *PALB2,* and 17 in *TP53*), probes were selected based on predicted T_m_s similar to other probes, limited alignment to other sequences in the transcriptome (< 10 times), and avoidance of stable, base-paired 2° structures (with unaFOLD) [69, 70]. The average coverage of these sequenced regions was 14.1–24.9% lower than other probe sets, indicating that capture was less efficient, though still successful.

### HBOC Samples for Oligo Capture and High-Throughput Sequencing

Genomic DNA used in prior susceptibility testing, from 102 anonymized patients was received from the Molecular Genetics Laboratory (MGL) at the London Health Sciences Centre in London, Ontario, Canada. Patients qualified for genetic susceptibility testing as determined by the Ontario Ministry of Health and Long-Term Care *BRCA1* and *BRCA2* genetic testing criteria [71] (see **Additional file 2**). *BRCA1* and *BRCA2* were previously analyzed by Protein Truncation Test (PTT) and Multiplex Ligation-dependent Probe Amplification (MLPA). The exons of several patients (N=14) had also been Sanger sequenced. No pathogenic sequence change was found in any of these individuals. In addition, one patient with a known pathogenic *BRCA* variant was re-sequenced by NGS as a positive control.

### Sequence Alignment and Variant Calling

Variant analysis involved the steps of detection, filtering, IT-based and coding sequence analysis, and prioritization (**Figure 2**). Sequencing data were demultiplexed and aligned to the specific chromosomes of our sequenced genes (hg19) using both CASAVA (Consensus Assessment of Sequencing and Variation; v1.8.2) [72] and CRAC (Complex Reads Analysis and Classification; v1.3.0) [73] software. Alignments were prepared for variant calling using Picard [74] and variant calling was performed on both versions of the aligned sequences using the UnifiedGenotyper tool in the Genome Analysis Toolkit (GATK) [75]. We used the recommended minimum phred base quality score of 30, and results were exported in variant call format (VCF; v4.1). A software program was developed to exclude variants called outside of targeted capture regions and those with quality scores < 50. Variants flagged by bioinformatic analysis (described below) were also assessed by manually inspecting the reads in the region using the Integrative Genomics Viewer (IGV; version 2.3) [76, 77] to note and eliminate obvious false positives (i.e. variant called due to polyhomonucleotide run dephasing, or PCR duplicates that were not eliminated by Picard). Finally, common variants (≥ 1% allele frequency based on dbSNP142 or > 5 individuals in our study cohort) were eliminated.

**Figure 2.**
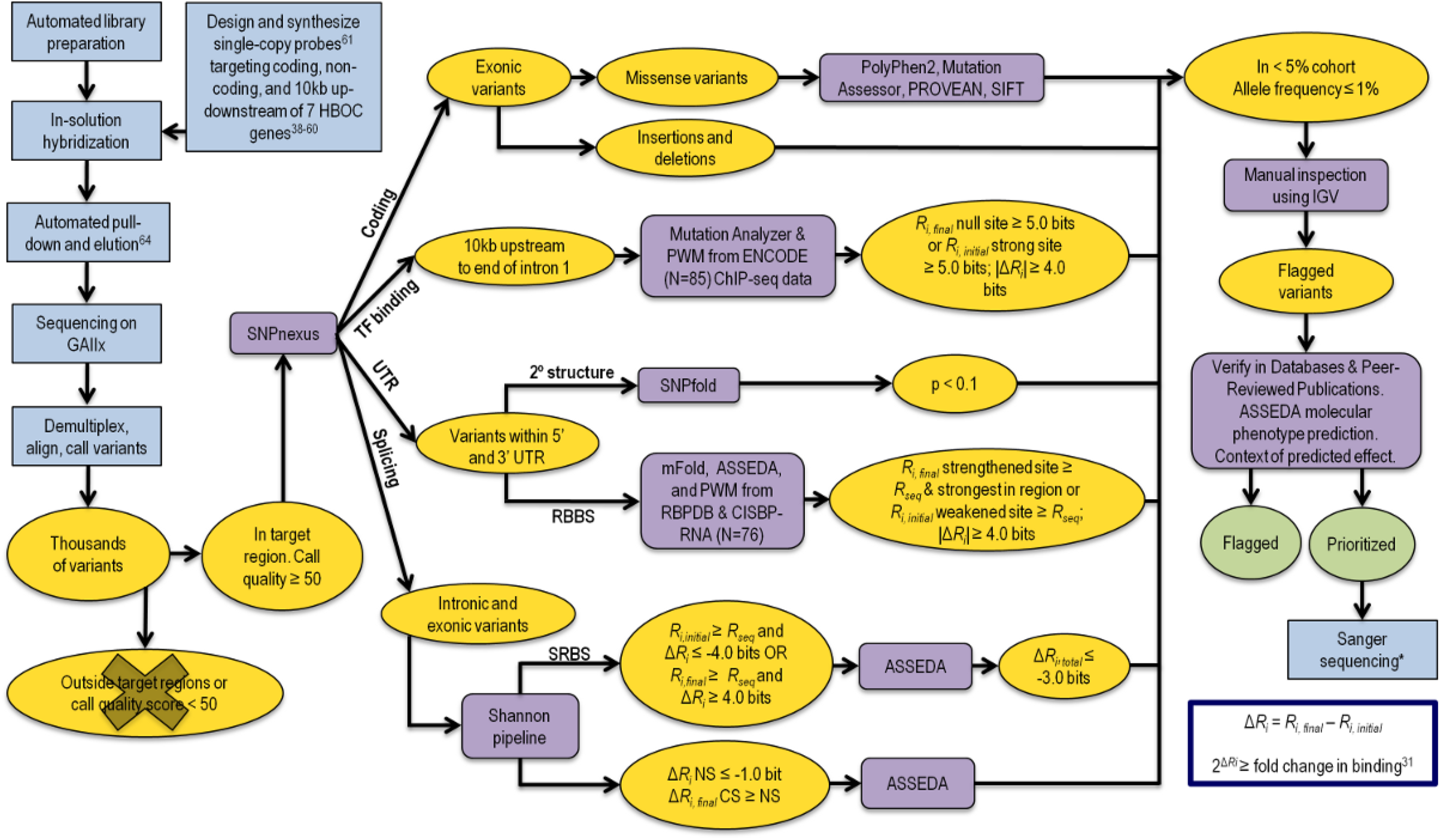
Framework for the Identification of Potentially Pathogenic Variants. Integrated laboratory processing and bioinformatic analysis procedures for comprehensive complete gene variant determination and analysis. Intermediate datasets resulting from filtering are represented in yellow and final datasets in green. Non-bioinformatic steps, such as sample preparation are represented in blue and prediction programs in purple. Sequencing analysis yields base calls for all samples. CASAVA [72] and CRAC [73] were used to align these sequencing results to HG19. GATK [75] was used to call variants from this data against GRCh37 release of the reference human genome. Variants with a quality score < 50 and/or call confidence score < 30 were eliminated along with variants falling outside of our target regions. SNPnexus [102–104] was used to identify the genomic location of the variants. Nonsense and indels were noted and prediction tools were used to assess the potential pathogenicity of missense variants. The Shannon Pipeline [78] evaluated the effect of a variant on natural and cryptic SSs, as well as SRFBSs. ASSEDA [80] was used to predict the potential isoforms as a result of these variants. PWMs for 83 TFs were built using an information weight matrix generator based on Bipad [95]. Mutation Analyzer evaluated the effect of variants found 10 kb upstream up to the first intron on protein binding. Bit thresholds (*R_i_* values) for filtering variants on software program outputs are indicated. Variants falling within the UTR sequences were assessed using SNPfold [15], and the most probable variants that alter mRNA structure (p < 0.1) were then processed using mFold to predict the effect on stability [70]. All UTR variants were scanned with a modified version of the Shannon Pipeline, which uses PWMs computed from nucleotide frequencies for 28 RBPs in RBPDB [99] and 76 RBPs in CISBP-RNA [100]. All variants meeting these filtering criteria were verified with IGV [76, 77]. Sanger sequencing was only performed for protein truncating, splicing, and selected missense variants

### IT-Based Variant Analysis

All variants were analyzed using the Shannon Human Splicing Mutation Pipeline, a genome-scale variant analysis program that predicts the effects of variants on mRNA splicing [78, 79]. Variants were flagged based on criteria reported in Shirley et al. (2013): weakened natural site ≥ 1.0 bits, or strengthened cryptic site (within 300 nt of the nearest exon) where cryptic site strength is equivalent or greater than the nearest natural site of the same phase [78]. The effects of flagged variants were further analyzed in detail using the Automated Splice Site and Exon Definition Analysis (ASSEDA) server [80].

Exonic variants and those found within 500 nt of an exon were assessed for their effects, if any, on SRFBSs [80]. Sequence logos for splicing regulatory factors (SRFs) (SRSF1, SRSF2, SRSF5, SRSF6, hnRNPH, hnRNPA1, ELAVL1, TIA1, and PTB) and their *R_sequence_* values (the mean information content [81]) are provided in Caminsky et al. (2015) [31]. Because these motifs occur frequently in unspliced transcripts, only variants with large information changes were flagged, notably those with (a) ≥ 4.0 bit decrease, i.e. at least a 16-fold reduction in binding site affinity, with *R_i,initial_* ≥ *R_sequence_* for the particular factor analyzed, or (b) ≥ 4.0 bit increase in a site where *R_i_,_final_* ≥ 0 bits. ASSEDA was used to calculate *R_i,total_,* with the option selected to include the given SRF in the calculation. Variants decreasing *R_i,total_* by < 3.0 bits (i.e. 8-fold) were predicted to potentially have benign effects on expression, and were not considered further.

Activation of pseudoexons through creating/strengthening of an intronic cryptic splice site was also assessed [82]. Changes in intronic cryptic sites, where ΔR*_i_* > 1 bit and *R_i_,_final_* ≥ (*R_sequence_* − 1 standard deviation [S.D.] of *R_sequence_*), were identified. A pseudoexon was predicted if a pre-existing cryptic site of opposite polarity (with *R_i_* > [*R_sequence_* − 1 S.D.]) and in the proper orientation for formation of exons between 10–250 nt in length was present. In addition, the minimum intronic distance between the pseudoexon and either adjacent natural exon was 100 nt. The acceptor site of the pseudoexon was also required to have a strong hnRNPA1 site located within 10 nt (*R_i_* ≥ *R_sequence_*) [80] to ensure accurate proofreading of the exon [83]. Next, variants affecting the strength of SRFs were analyzed by a contextual exon definition analysis of Δ*R_i,total_.* The context refers to the documented splicing activity of an SRF. For example, TIA1 has been shown to be an intronic enhancer of exon definition, so only intronic sites were considered. Similarly, hnRNPA1 proofreads the 3′ SS (acceptor) and inhibits exon recognition elsewhere [84]. Variants that lead to redundant SRFBS changes (i.e. one site is abolished and another proximate site [≤ 2 nt] of equivalent strength is activated) were assumed to have a neutral effect on splicing. If the strength of a site bound by PTB (polypyrimidine tract binding protein) was affected, its impact on binding by other factors was analyzed, as PTB impedes binding of other factors with overlapping recognition sites, but does not directly enhance or inhibit splicing itself [85].

To determine effects of variants on transcription factor (TF) binding, we first established which TFs bound to the sequenced regions of the gene promoters (and first exons) in this study by using ChIP-seq data from 125 cell types (**Supplementary Methods**) [86]. We identified 141 TFs with evidence for binding to the promoters of the genes we sequenced, including c-Myc, C/EBPβ, and Sp1, shown to transcriptionally regulate *BRCA1, TP53,* and *ATM,* respectively [87–89]. Furthermore, polymorphisms in TCF7L2, known to bind enhancer regions of a wide variety of genes in a tissue-specific manner [90], have been shown to increase risk of sporadic [91] and hereditary breast [92], as well as other types of cancer [93, 94].

IT-based models of the 141 TFs of interest were derived by entropy minimization of the DNase accessible ChIP-seq subsets [95]. Details are provided in another concurrently submitted manuscript (Liu et al. submitted: included for purposes of review). While some data sets would only yield noise or co-factor motifs (i.e. co-factors that bind via tethering, or histone modifying proteins [96]), techniques such as motif masking and increasing the number of Monte Carlo cycles yielded models for 83 TFs resembling each factor’s published motif. **Table S1** (**Additional file 3**) contains the final list of TFs and the models we built (described below) [97].

These TFBS models (N=83) were used to scan all variants called in the promoter regions (10 kb upstream of transcriptional start site to the end of IVS1) of HBOC genes for changes in *R_i_* [98] Binding site changes that weaken interactions with the corresponding TF (to *R_i_* ≤ *R_sequence_*) are likely to affect regulation of the adjacent target gene. Stringent criteria were used to prioritize the most likely variants and thus only changes to strong TFBSs (*R_i_*,*_initial_* ≥ *R_sequence_*), where reduction in strength was significant (Δ*R_i_* ≥ 4.0 bits), were considered. Alternatively, novel or strengthened TFBSs were also considered sources of dysregulated transcription. These sites were defined as having *R_i_,_final_* ≥ *R_sequence_* and as being the strongest predicted site in the corresponding genomic interval (i.e. exceeding the *R_i_* values of adjacent sites unaltered by the variant). Variants were not prioritized if the TF was known to a) enhance transcription and IT analysis predicted stronger binding, or b) repress transcription and IT analysis predicted weaker binding.

Two complementary strategies were used to assess the possible impact of variants within UTRs. First, SNPfold software was used to assess the effect of a variant on 2° structure of the UTR (**Supplementary Methods**) [15]. Variants flagged by SNPfold with the highest probability of altering stable 2° structures in mRNA (where p-value < 0.1) were prioritized. To evaluate these predictions, oligonucleotides containing complete wild-type and variant UTR sequences (**Table S2 – Additional file 4**) were transcribed *in vitro* and followed by SHAPE analysis, a method that can confirm structural changes in mRNA [35].

Second, the effects of variants on the strength of RBBSs were predicted. Frequency-based, position weight matrices (PWMs) for 156 RNA-binding proteins (RBPs) were obtained from the RNA-Binding Protein DataBase (RBPDB) [99] and the Catalog of Inferred Sequence Binding Preferences of RNA binding proteins (CISBP-RNA) [100, 101]. These were used to compute information weight matrices (based on the method described by Schneider et al. 1984; N = 147) (see **Supplementary Methods**) [32]. All UTR variants were assessed using a modified version of the Shannon Pipeline [78] containing the RBPDB and CISBP-RNA models. Results were filtered to include a) variants with |Δ*R_i_*| ≥ 4.0 bits, b) variants creating or strengthening sites (*R_i.final_* ≥ *R_sequence_* and the *R_i,initial_* < *R_sequence_*), and c) RBBSs not overlapping or occurring within 10 nt of a stronger, pre-existing site of another RBP.

### Exonic Protein-Altering Variant Analysis

The predicted effects of all coding variants were assessed with SNPnexus [102–104], an annotation tool that can be applied to known and novel variants using up-to-date dbSNP and UCSC human genome annotations. Variants predicted to cause premature protein truncation were given higher priority than those resulting in missense (or synonymous) coding changes. Missense variants were first cross referenced with dbSNP142 [105]. Population frequencies from the Exome Variant Server [106] and 1000Genomes [107] are also provided. The predicted effects on protein conservation and function of the remaining variants were evaluated by *in silico* tools: PolyPhen-2 [108], Mutation Assessor (release 2) [109, 110], and PROVEAN (v1.1.3) [111, 112]. Default settings were applied and in the case of PROVEAN, the “PROVEAN Human Genome Variants Tool” was used, which includes SIFT predictions as a part of its output. Variants predicted by all four programs to be benign were less likely to have a deleterious impact on protein activity; however this did not exclude them from mRNA splicing analysis (described above in *IT-Based Variant Analysis*). All rare and novel variants were cross-referenced with general mutation databases (ClinVar [113, 114], Human Gene Mutation Database [HGMD] [115, 116], Leiden Open Variant Database [LOVD] [117–124], Domain Mapping of Disease Mutations [DM^2^] [125], Expert Protein Analysis System [ExPASy] [126] and UniProt [127, 128]), and gene-specific databases (*BRCA1/2:* the Breast Cancer Information Core database [BIC] [129] and Evidence-based Network for the Interpretation of Germline Mutant Alleles [ENIGMA] [130]; *TP53:* International Agency for Research on Cancer [IARC] [131]), as well as published reports to prioritize them for further workup.

### Variant Classification

Flagged variants were prioritized if they were likely to encode a dysfunctional protein (indels, nonsense codon > 50 amino acids from the C-terminus, or abolition of a natural SS resulting in out-of-frame exon skipping) or if they exceeded established thresholds for fold changes in binding affinity based on IT (see *Methods* above). If previous studies performed functional or pedigree analyses, allowing to categorize a variant as pathogenic or benign, this superseded our analysis.

### Positive control

We identified the *BRCA1* exon 17 nonsense variant c.5136G>A (chr17:41215907C>T; rs80357418; 2–5A) [132] in the sample that was provided as a positive control. This was the same mutation identified by the MGL as pathogenic for this patient. We also prioritized another variant in this patient (**Table 1**) [133].

**Table 1.** Prioritized Variants in the Positive Control.

| Gene | mRNA<br>Protein | rsID (dbSNP142)<br>Allele Frequency (%)<sup>†</sup> | Category | Consequence | Ref |
| --- | --- | --- | --- | --- | --- |
| *BRCA1* | c.5136G>A<br>p.Trp1712Ter | rs80357418 | Nonsense | 151 AA short | [132] |
| *BRCA2* | c.3218A>G<br>p.Gln1073Arg | rs80358566 | Missense | Listed in ClinVar as conflicting interpretations (likely benign, unknown) and in BIC as unknown clinical importance. 2 *in silico* programs called deleterious. | [133] |
|  |  |  | SRFBS | Repressor action of hnRNPA1 at this site abolished (5.2 to 0.4 bits). Blocking action of PTB removed as site is abolished (5.5 to -7.5 bits) and may uncover binding sites of other SRFs. |  |

<sup>†</sup> If available. Positive control was sample 2-5A.

### Variant Validation

Protein-truncating, prioritized splicing, and selected prioritized missense variants were verified by Sanger sequencing. Primers of PCR amplicons are indicated in **Table S3 – Additional File 5**).

## RESULTS

### Capture, Sequencing, and Alignment

The average coverage of capture region per individual was 90.8x (range of 53.8 to 118.2× between 32 samples) with 98.8% of the probe-covered nucleotides having ≥ 10 reads. Samples with fewer than 10 reads per nucleotide were re-sequenced and the results of both runs were combined. The combined coverage of these samples was, on average, 48.2× (± 36.2).

The consistency of both library preparation and capture protocols was improved from initial runs, which significantly impacted sequence coverage (**Supplementary Methods**). Of the 102 patients tested, 14 had been previously Sanger sequenced for *BRCA1* and *BRCA2* exons. Confirmation of previously discovered SNVs served to assess the methodological improvements introduced during NGS and ultimately, to increase confidence in variant calling. Initially, only 15 of 49 SNVs in 3 samples were detected. The detection rate of SNVs was improved to 100% as the protocol progressed. All known SNVs (N=157) were called in subsequent sequencing runs where purification steps were replaced with solid phase reversible immobilization beads and where RNA bait was transcribed the same day as capture. To minimize false positive variant calls, sequence read data was aligned using 2 different software programs, CASAVA and CRAC, and variant calling was performed for both sets of data using GATK [72, 73, 75].

GATK called 14,164 unique SNVs and 1,147 indels. Only 3,777 (15.3%) SNVs were present in both CASAVA and CRAC-alignments for at least one patient, and even fewer indel calls were concordant between both methods (N=110; 6.2%). For all other SNVs and indels, CASAVA called 6,871 and 1,566, respectively, whereas CRAC called 13,958 and 110, respectively. Some variants were counted more than once if they are called by different alignment programs in two or more patients. Intronic and intergenic variants proximate to low complexity sequences tend to generate false positive variants due to ambiguous alignment, a well known technical issue in short read sequence analysis [134, 135], contributing to this discrepancy. For example, in **Figure S1** (**Additional file 6**), CRAC correctly called a 19 nt deletion of *BRCA1* (rs80359876; also confirmed by Sanger sequencing) but CASAVA flagged the deleted segment as a series of false-positives. For these reasons, all variants were manually reviewed.

### IT-Based Variant Identification and Prioritization

#### Natural SS Variants

The Shannon Pipeline reported 99 unique variants in natural donor or acceptor SSs. After technical and frequency filtering criteria were applied, 12 variants remained (**Table S4 - Additional file 7**). IT analysis allowed for the prioritization of 3 variants, summarized in **Table 2**.

**Table 2.**
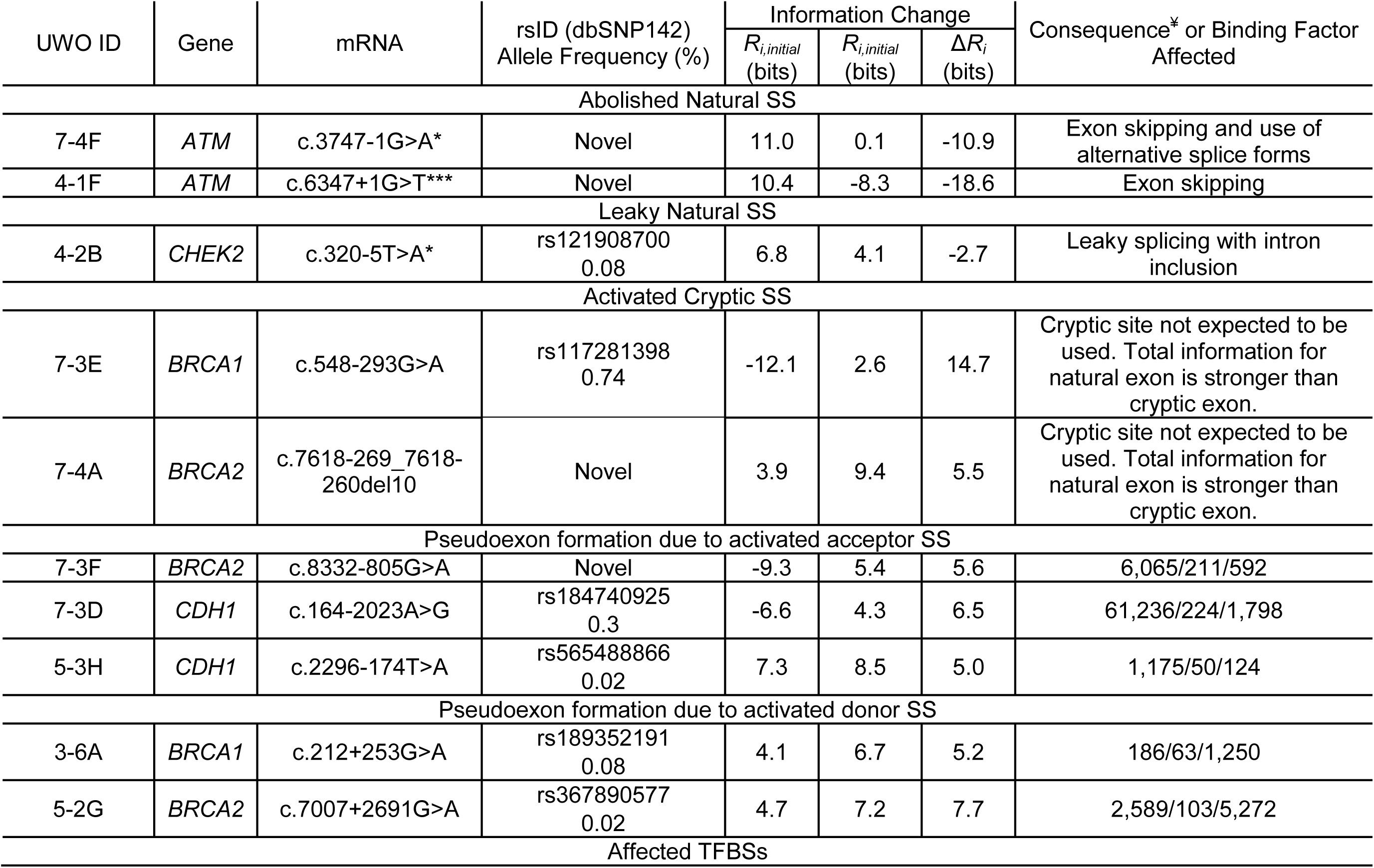
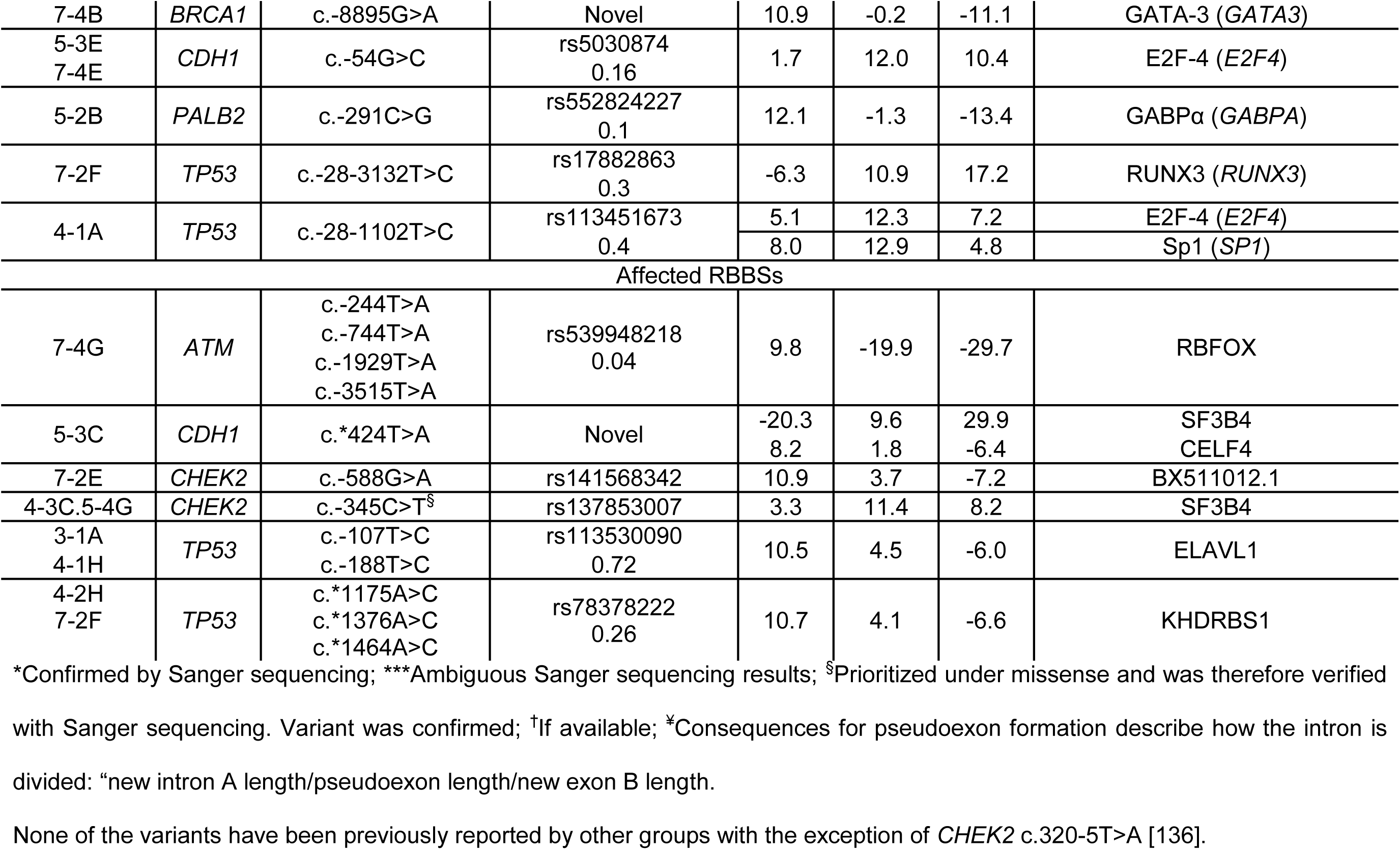
Variants Prioritized by IT Analysis.

| UWO ID | Gene | mRNA | rsID (dbSNP142)<br>Allele Frequency (%) | Information Change |  |  | Consequence* or Binding Factor Affected |
| --- | --- | --- | --- | --- | --- | --- | --- |
| | | | | $R_{i,initial}$<br>(bits) | $R_{i,initial}$<br>(bits) | $\Delta R_i$<br>(bits) | |
| Abolished Natural SS |  |  |  |  |  |  |  |
| 7-4F | ATM | c.3747-1G>A* | Novel | 11.0 | 0.1 | -10.9 | Exon skipping and use of alternative splice forms |
| 4-1F | ATM | c.6347+1G>T*** | Novel | 10.4 | -8.3 | -18.6 | Exon skipping |
| Leaky Natural SS |  |  |  |  |  |  |  |
| 4-2B | CHEK2 | c.320-5T>A* | rs121908700<br>0.08 | 6.8 | 4.1 | -2.7 | Leaky splicing with intron inclusion |
| Activated Cryptic SS |  |  |  |  |  |  |  |
| 7-3E | BRCA1 | c.548-293G>A | rs117281398<br>0.74 | -12.1 | 2.6 | 14.7 | Cryptic site not expected to be used. Total information for natural exon is stronger than cryptic exon. |
| 7-4A | BRCA2 | c.7618-269_7618-260del10 | Novel | 3.9 | 9.4 | 5.5 | Cryptic site not expected to be used. Total information for natural exon is stronger than cryptic exon. |
| Pseudoexon formation due to activated acceptor SS |  |  |  |  |  |  |  |
| 7-3F | BRCA2 | c.8332-805G>A | Novel | -9.3 | 5.4 | 5.6 | 6,065/211/592 |
| 7-3D | CDH1 | c.164-2023A>G | rs184740925<br>0.3 | -6.6 | 4.3 | 6.5 | 61,236/224/1,798 |
| 5-3H | CDH1 | c.2296-174T>A | rs565488866<br>0.02 | 7.3 | 8.5 | 5.0 | 1,175/50/124 |
| Pseudoexon formation due to activated donor SS |  |  |  |  |  |  |  |
| 3-6A | BRCA1 | c.212+253G>A | rs189352191<br>0.08 | 4.1 | 6.7 | 5.2 | 186/63/1,250 |
| 5-2G | BRCA2 | c.7007+2691G>A | rs367890577<br>0.02 | 4.7 | 7.2 | 7.7 | 2,589/103/5,272 |
| Affected TFBSs |  |  |  |  |  |  |  |
| 7-4B | *BRCA1* | c.-8895G>A | Novel | 10.9 | -0.2 | -11.1 | GATA-3 (*GATA3*) |
| 5-3E<br>7-4E | *CDH1* | c.-54G>C | rs5030874<br>0.16 | 1.7 | 12.0 | 10.4 | E2F-4 (*E2F4*) |
| 5-2B | *PALB2* | c.-291C>G | rs552824227<br>0.1 | 12.1 | -1.3 | -13.4 | GABPα (*GABPA*) |
| 7-2F | *TP53* | c.-28-3132T>C | rs17882863<br>0.3 | -6.3 | 10.9 | 17.2 | RUNX3 (*RUNX3*) |
| 4-1A | *TP53* | c.-28-1102T>C | rs113451673<br>0.4 | 5.1 | 12.3 | 7.2 | E2F-4 (*E2F4*) |
|  |  |  |  | 8.0 | 12.9 | 4.8 | Sp1 (*SP1*) |
| Affected RBBs |  |  |  |  |  |  |  |
| 7-4G | *ATM* | c.-244T>A<br>c.-744T>A<br>c.-1929T>A<br>c.-3515T>A | rs539948218<br>0.04 | 9.8 | -19.9 | -29.7 | RBFOX |
| 5-3C | *CDH1* | c.*424T>A | Novel | -20.3<br>8.2 | 9.6<br>1.8 | 29.9<br>-6.4 | SF3B4<br>CELF4 |
| 7-2E | *CHEK2* | c.-588G>A | rs141568342 | 10.9 | 3.7 | -7.2 | BX511012.1 |
| 4-3C.5-4G | *CHEK2* | c.-345C>T<sup>§</sup> | rs137853007 | 3.3 | 11.4 | 8.2 | SF3B4 |
| 3-1A<br>4-1H | *TP53* | c.-107T>C<br>c.-188T>C | rs113530090<br>0.72 | 10.5 | 4.5 | -6.0 | ELAVL1 |
| 4-2H<br>7-2F | *TP53* | c.*1175A>C<br>c.*1376A>C<br>c.*1464A>C | rs78378222<br>0.26 | 10.7 | 4.1 | -6.6 | KHDRBS1 |

\*Confirmed by Sanger sequencing; \*\*\*Ambiguous Sanger sequencing results; <sup>§</sup>Prioritized under missense and was therefore verified with Sanger sequencing. Variant was confirmed; <sup>†</sup>If available; \*Consequences for pseudoexon formation describe how the intron is divided: "new intron A length/pseudoexon length/new exon B length.
None of the variants have been previously reported by other groups with the exception of *CHEK2* c.320-5T>A [136].

First, the novel *ATM* variant c.3747–1G>A (chr11:108154953G>A; sample number 7–4F) abolishes the natural acceptor of exon 26 (11.0 to 0.1 bits). ASSEDA reports the presence of a 5.3 bit cryptic acceptor site 13 nt downstream of the natural site, but the effect of the variant on a pre-existing cryptic site is negligible (~0.1 bits). The cryptic exon would lead to exon deletion and frameshift (**Figure 3A**). ASSEDA also predicts skipping of the 246 nt exon, as the *R_i,final_* of the natural acceptor is now below *R_i_,_minimum_* (16 bits), altering the reading frame. Second, the novel *ATM* c.6347+1G>T (chr11:108188249G>T; 4–1F) occurs at the natural donor of exon 44 and abolishes the 10.4 bit donor (Δ*R_i_* = −18.6 bits), resulting exclusively in exon skipping. Finally, the previously reported *CHEK2* variant, c.320–5A>T (chr22:29121360T>A; rs121908700; 4–2B) [136] weakens the natural acceptor of exon 3 (6.8 to 4.1 bits), possibly activating a cryptic acceptor (7.4 bits) 92 nt upstream of the natural acceptor (**Figure 4**).

**Figure 3.**
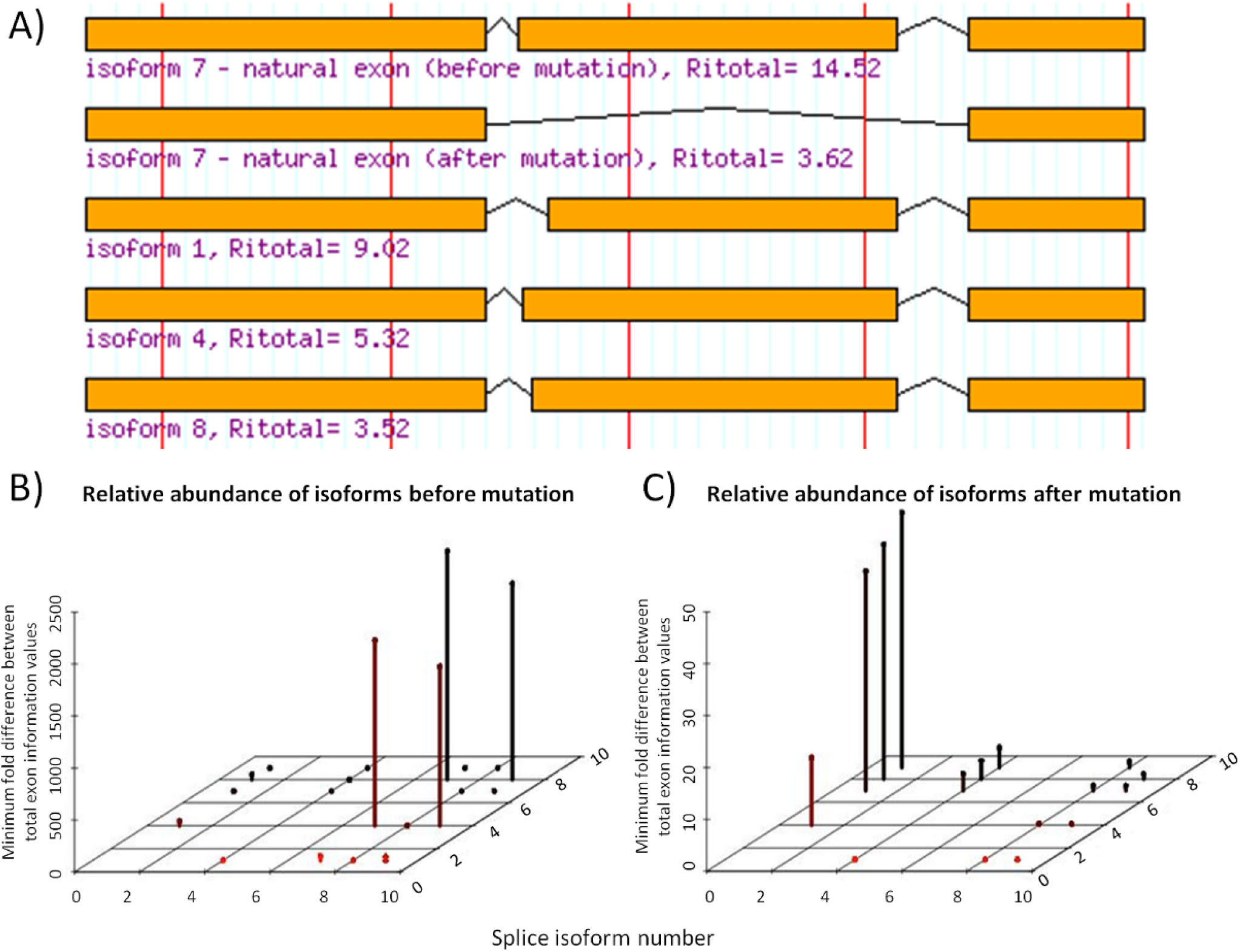
Predicted Isoforms and Relative Abundances as a Consequence of *ATM* splice variant c.3747–1G>A. Intronic *ATM* variant c.3747–1G>A abolishes (11.0 to 0.1 bits) the natural acceptor of exon 26 (total of 63 exons). **A)** ASSEDA reports the abolition of the natural exon (*R_i,total_* reduced from 14.5 to 3.6 bits) and predicts exon skipping as a result (isoform 7 after mutation) and/or the use of a cryptic site 13 nt downstream (*R_i,total_* for cryptic exon = 9.0 bits) of the natural site leading to exon deletion (isoform 1). The other isoforms use weak, alternate acceptor/donor sites leading to cryptic exons with much lower total information. **B)** Before the mutation, isoform 7 is expected to be the most abundant splice form. **C)** After the mutation, isoform 1 is predicted to become the most abundant splice form and the wild-type isoform is not expected to be expressed.

**Figure 4.**
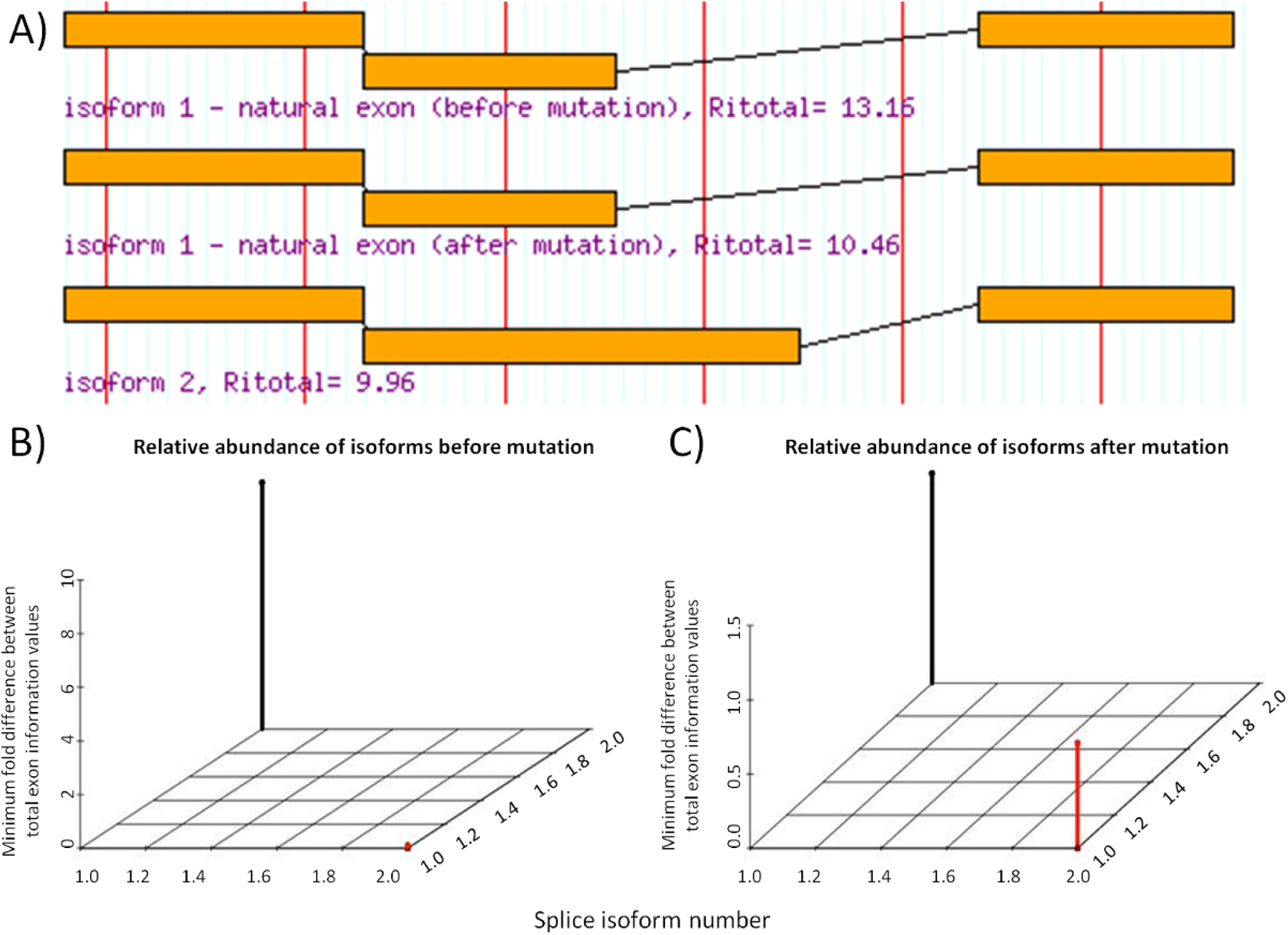
Predicted Isoforms and Relative Abundances as a Consequence of *CHEK2* splice variant c.320–5T>A. Intronic *CHEK2* variant c.320–5T>A weakens (6.8 to 4.1 bits) the natural acceptor of exon 3 (total of 15 exons). **A)** ASSEDA reports the weakening of the natural exon strength (*R_i,total_* reduced from 13.2 to 10.5 bits), which would result in reduced splicing of the exon otherwise known as leaky splicing. A pre-existing cryptic acceptor exists 92 nt upstream of the natural site, leading to a cryptic exon with similar strength to the mutated exon (*R_i,total_* = 10.0 bits). This cryptic exon would contain 92 nt of the intron. **B)** Before the mutation, isoform 1 is expected to be the only isoform expressed. **C)** After the mutation, isoform 1 (wild-type) is predicted to become relatively less abundant and isoform 2 is expected to be expressed, although less abundant in relation to isoform 1.

Variants either strengthening (N=4) or slightly weakening (Δ*R_i_* < 1.0 bits; N=4) a natural site were not prioritized. In addition, we rejected the *ATM* variant (c.1066–6T>G; chr11:108119654T>G; 4–1E and 7–2B), which slightly weakens the natural acceptor of exon 9 (11.0 to 8.1 bits). Although other studies have shown leaky expression as a result of this variant [137], a more recent meta-analysis concluded that this variant is not associated with increased breast cancer risk [138].

#### Cryptic SS Activation

Two variants produced information changes that could potentially impact cryptic splicing, but were not prioritized for the following reasons (**Table 2**). The first variant, novel *BRCA2* deletion c.7618–269_7618–260del10 (chr13:32931610_32931619del10; 7–4A) strengthens a cryptic acceptor site 245 nt upstream from the natural acceptor of exon 16 (*R_i,final_* = 9.4 bits, Δ*R_i_* = 5.5 bits). Being 5.7-fold stronger than the natural site (6.9 bits), two potential cryptic isoforms were predicted, however, the exon strengths of both are weaker than the unaffected natural exon (*R_i_,_total_* = 6.6 bits) and neither were prioritized. The larger gap surprisal penalties explain the differences in exon strength. The natural donor SS may still be used in conjunction with the abovementioned cryptic SS, resulting in an exon with *R_i,total_* = 3.5 bits. Alternatively, the cryptic site and a weak donor site 180 nt upstream of the natural donor (*R_i_* = 0.7 vs 1.4, cryptic and natural donors, respectively), result in an exon with *R_i,total_* = 6.5 bits. The second variant, *BRCA1* c.548–293G>A (chr17:41249599C>T; 7–3E), creates a weak cryptic acceptor (*R_i,final_* = 2.6 bits, Δ*R_i_* = 6.2 bits) 291 nt upstream of the natural acceptor for exon 8 (*R_i_* = 0.5). Although the cryptic exon is strengthened (final *R_i,total_* = 6.9 bits, Δ*R_i_* = 14.7 bits), ASSEDA predicts the level of expression of this exon to be negligible, as it is weaker than the natural exon (*R_i,total_* = 8.4 bits) due to the increased length of the predicted exon (+291 nt) [80].

#### Pseudoexon Formation

The Shannon Pipeline initially reported 1,583 unique variants creating or strengthening intronic cryptic sites. We prioritized 5 variants, 1 of which is novel (*BRCA2* c.8332–805G>A; 7–3F), that were within 250 nt of a pre-existing complementary cryptic site and have an hnRNPA1 site within 5 nt of the acceptor (**Table 2**). If used, 3 of these pseudoexons would lead to a frameshifted transcript.

#### SRF Binding

Variants within 500 nt of an exon junction and all exonic variants (N = 4,015) were investigated for their potential effects on affinity of sites to corresponding SRFs [80]. IT analysis flagged 54 variants significantly altering the strength of at least one binding site (**Table S5 - Additional file 8**). A careful review of the variants, the factor affected, and the position of the binding site relative to the natural SS, prioritized 36 variants (21 novel), of which 4 are in exons and 32 are in introns.

#### TF Binding

We assessed SNVs with models of 83 TFs experimentally shown to bind (**Table S1**) upstream or within the first exon and intron of our sequenced genes (N=2,177). Thirteen variants expected to significantly affect TF binding were flagged (**Table S6 - Additional file 9**). The final filtering step considered the known function of the TF in transcription, resulting in 5 prioritized variants (**Table 2**) in 6 patients (one variant was identified in two patients). Four of these variants have been previously reported (rs5030874, rs552824227, rs17882863, rs113451673) and one is novel (c.-8895G>A; 7–4B).

#### UTR Structure and Protein Binding

There were 364 unique UTR variants found by sequencing, which includes splice forms with alternate UTRs (in *BRCA1* and *TP53*). These variants were evaluated for their effects on mRNA 2° structure through SNPfold, resulting in 5 flagged variants (**Table 3**), all of which have been previously reported.

**Table 3.** Variants Predicted by SNPfold to Affect UTR Structure.

| Class<sup>¥</sup> | UWO ID | Gene | mRNA | UTR position | rsID (dbSNP142)<br>Allele Frequency (%)<sup>†</sup> | Rank<sup>§</sup> | p-value |
| --- | --- | --- | --- | --- | --- | --- | --- |
| F | In 26 patients | *BRCA2*<sup>§</sup> | c.-52A>G | 5' UTR | rs206118<br>14.86 | 2/900 | 0.002 |
| F | In 40 patients | *BRCA2*<sup>§</sup> | c.*532A>G | 3' UTR | rs11571836<br>19.75 | 239/2700 | 0.089 |
| P | 7-4C | *CDH1*<sup>§§</sup> | c.-71C>G | 5' UTR | rs34033771<br>0.56 | 69/600 | 0.115 |
| F | 4-2E<br>5-4A | *TP53*<sup>§</sup> | c.*485G>A | 3' UTR | rs4968187<br>5.11 | 169/4500 | 0.038 |
| F | 2-1A, 7-1B,<br>5-2A.7-1D,<br>7-2B, 7-2F<br>7-4C | *TP53*<sup>§</sup> | c.*826G>A | 3' UTR | rs17884306<br>5.71 | 371/4500 | 0.082 |

<sup>¥</sup>F:Flagged; P:Prioritized; <sup>§</sup>Long Range UTR SNPfold Analysis; <sup>§§</sup>Local Range SNPfold Analysis; <sup>†</sup>If available; <sup>§</sup>Rank of the SNP, in terms of how much it changes the mRNA structure compared to all other possible mutations.

Analysis of three variants using mFOLD [70] revealed likely changes to the UTR structure (**Figure 5**). Two variants with possible 2° structure effects were common (*BRCA2* c.-52A>G [N=26 samples] and c.*532A>G [N=40]) and not prioritized. The 5’UTR *CDH1* variant c.-71C>G (chr16:68771248C>G; rs34033771; 7–4C) disrupts a double-stranded hairpin region to create a larger loop structure, thus increasing binding accessibility (**Figure 5A** and **B**). Analysis using RBPDB and CISBP-RNA-derived IT models suggests this variant affects binding by NCL by decreasing binding affinity 14-fold (R*_i_*,*_initial_* = 6.6 bits, Δ*R_i_* = −3.8 bits) (**Table S7** - **Additional file 10**). This RBP has been shown to bind to the 5’ and 3’ UTR of p53 mRNA and plays a role in repressing its translation [139].

**Figure 5.**
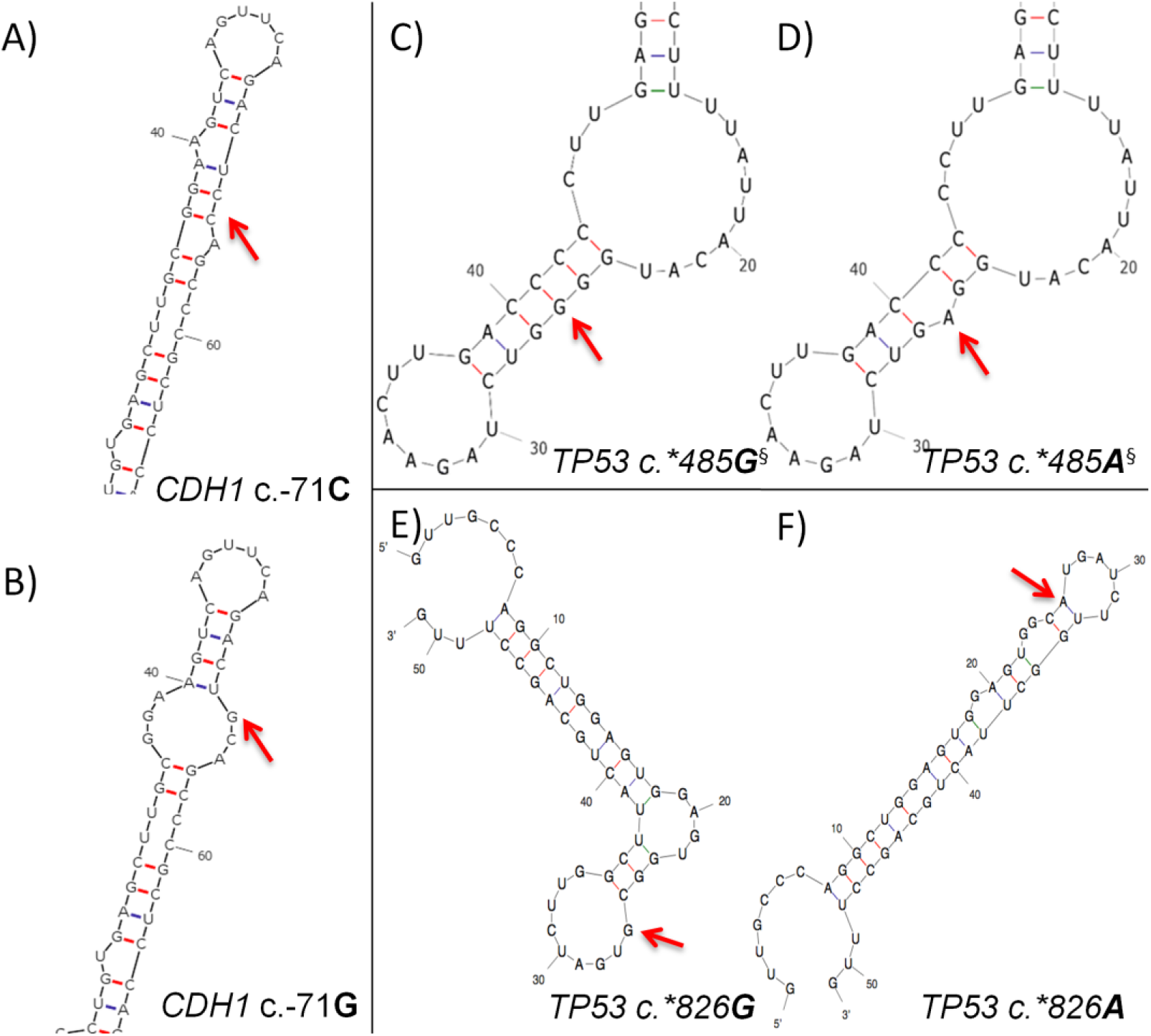
Predicted Alteration in UTR Structure Using mFOLD for Variants Flagged by SNPfold. Wild-type and variant structures are displayed, with the variant indicated by a red arrow. **A)** Predicted wild-type structure of *CDH1* 5’UTR surrounding c.-71. **B)** Predicted *CDH1* 5’UTR structure due to c.-71C>G variant. **C)** Predicted wild-type *TP53* 3’UTR structure surrounding c.*485. **D)** Predicted *TP53* 5’UTR structure due to c.*485G>A variant. **E)** Predicted wild-type *TP53* 3’UTR structure surrounding c.*826. **F)** Predicted *TP53* 5’UTR structure due to c.*826G>A variant. ^§^SHAPE analysis revealed differences in reactivity between mutant and variant mRNAs, confirming alterations to 2° structure.

In addition, the *TP53* variant c.*485G>A (NM_000546.5: chr17:7572442C>T; rs4968187) is found at the 3’UTR and was identified in two patients (4–2E and 5–4A). *In silico* mRNA folding analysis demonstrates this variant disrupts a G/C bond of a loop in the highest ranked potential mRNA structure (**Figure 5C and D**). Also, SHAPE analysis shows a difference in 2° structure between the wild-type and mutant (data not shown). IT analysis with RBBS models indicated that this variant significantly increases the binding affinity of SF3B4 > 48-fold (*R_i,final_* = 11.0 bits, Δ*R_i_* = 5.6 bits) (**Table S7**). This RBP is one of four subunits comprising the splice factor 3B and is known to bind upstream of the branch-point sequence in pre-mRNA [140].

The third flagged variant also occurs in the 3’UTR of *TP53* (c.*826G>A; chr17:7572101C>T; rs17884306), and was identified in 6 patients (2–1A, 7–1B, 5–2A.7–1D, 7–2B, 7–2F, and 7–4C). It disrupts a potential loop structure, stabilizing a double-stranded hairpin, and possibly making it less accessible (**Figure 5E and F**). Analysis using RBPDB-derived models suggests this variant could affect the binding of both RBFOX2 and SF3B4 (**Table S7**). A binding site for RBFOX2, which acts as a promoter of alternative splicing by favoring the inclusion of alternative exons [141], is created (*R_i,final_* = 9.8 bits; Δ*R_i_* = −6.5 bits). This variant is also expected to simultaneously abolish a SF3B4 binding site (*R_i,final_* = −20.3 bits; Δ*R_i_* = −29.9 bits).

RBPDB and CISBP-RNA-derived information model analysis of all UTR variants resulted in the prioritization of 1 novel, and 5 previously-reported variants (**Table 2**). No patient within the cohort exhibits more than one prioritized RBBS variant.

### Exonic Variants altering protein sequence

Exonic variants called by GATK (N=245) included insertions, deletions, nonsense, missense, and synonymous changes.

#### Protein-Truncating Variants

We identified 3 patients with different indels (**Table 4**). One was a *PALB2* insertion c.1617_1618insTT (chr16:23646249_23646250insAA; 5–3A) in exon 4, previously reported in ClinVar as pathogenic. This mutation results in a frameshift and premature translation termination by 626 residues, abolishing domain interactions with RAD51, BRCA2, and POLH [127]. We also identified two known frameshift mutations in *BRCA1:* c.4964_4982del19 in exon 15 (chr17:41222949_41222967del19; rs80359876; 5–1B) and c.5266_5267insC in exon 19 (chr17:41209079_41209080insG; rs397507247; 5–3C) [136, 142]. Both are indicated as pathogenic and common in the BIC Database due to the loss of one or both C-terminal BRCT repeat domains [127]. Truncation of these domains produces instability and impairs nuclear transcript localization [143], and this bipartite domain is responsible for binding phosphoproteins that are phosphorylated in response to DNA damage [144, 145].

**Table 4.**
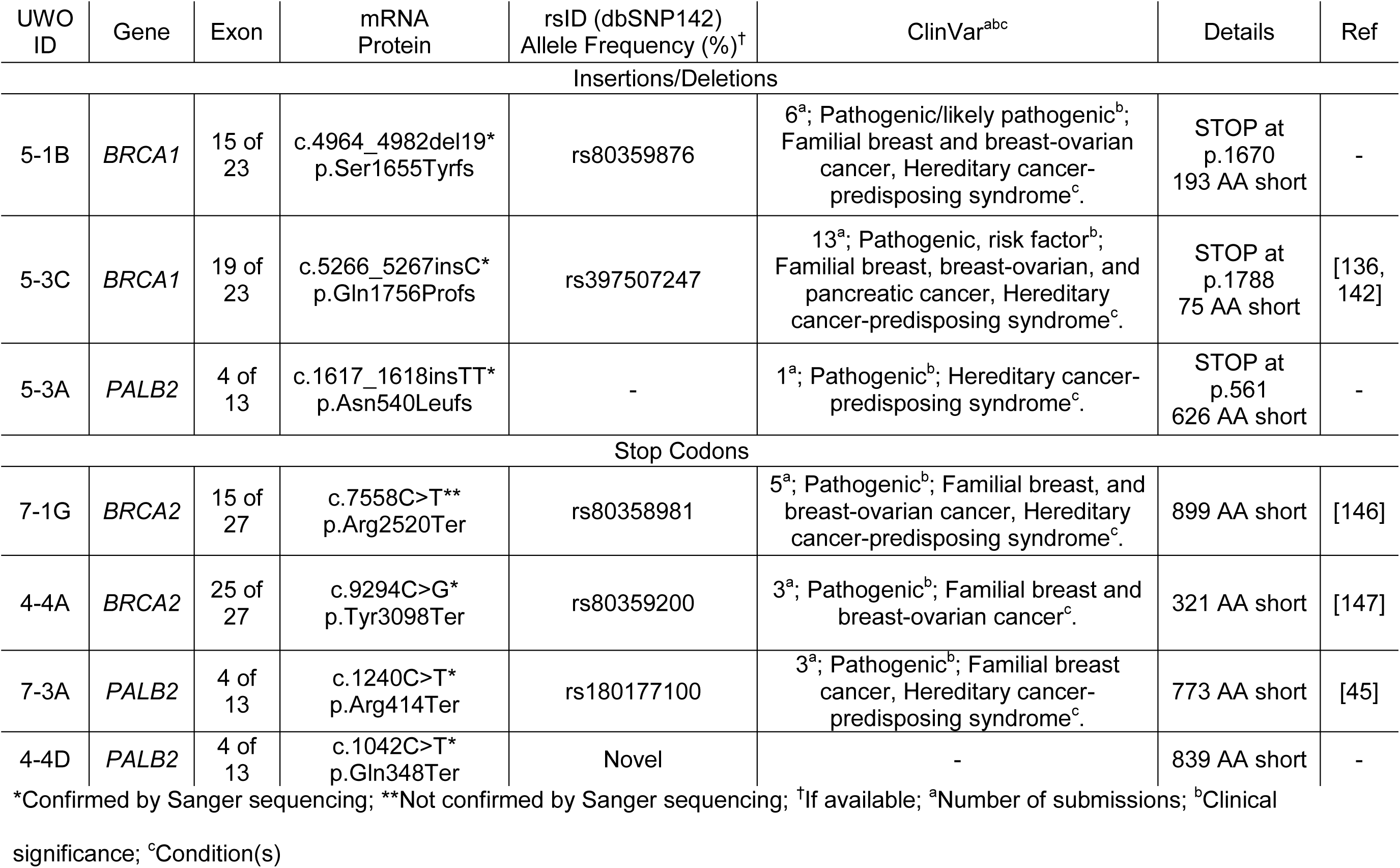
Variants Resulting in Premature Protein Truncation.

We also identified 4 nonsense mutations, one of which was novel in exon 4 of *PALB2* (c.1042C>T; chr16:23646825G>A; 4–4D). Another in *PALB2* has been previously reported (c.1240C>T; chr16:23646627G>A; rs180177100; 7–3A) [45]. As a consequence, functional domains of PALB2 that interact with BRCA1, RAD51, BRCA2, and POLH are lost [127]. Two known nonsense mutations were found in *BRCA2,* c.7558C>T in exon 15 [146] and c.9294C>G in exon 25 [147]. The first (chr13:32930687C>T; rs80358981; 7–1G) causes the loss of the BRCA2 region that binds FANCD2, which loads BRCA2 onto damaged chromatin [148]. The second (chr13:32968863C>G, rs80359200; 4–4A) does not occur within a known functional domain, however the transcript is likely to be degraded by nonsense mediated decay [149].

#### Missense

GATK called 61 missense variants, of which 18 were identified in 6 patients or more and 19 had allele frequencies > 1.0% (**Table S8 - Additional file 11**). The 40 remaining variants (15 *ATM,* 8 *BRCA1,* 9 *BRCA2,* 2 *CDH1,* 2 *CHEK2,* 3 *PALB2,* and 1 *TP53*) were assessed using a combination of gene specific databases, published classifications, and 4 *in silico* tools (**Table S9 - Additional file 12**). We prioritized 27 variants, 2 of which were novel. None of the non-prioritized variants were predicted to be damaging by more than 2 of 4 conservation-based software programs.

### Variant Classification

Initially, 15,311 unique variants were identified by complete gene sequencing of 7 HBOC genes. Of these, 132 were flagged after filtering, and further reduced by IT-based variant analysis and consultation of the published literature to 87 prioritized variants. **Figure 6** illustrates the decrease in the number of unique variants per patient at each step of our identification and prioritization process. The distribution of prioritized variants by gene is 34 in *ATM,* 13 in *BRCA1,* 11 in *BRCA2,* 8 in *CDH1,* 6 in *CHEK2,* 10 in *PALB2,* and 5 in *TP53* (**Table S10 - Additional file 13**), which are categorized by type in **Table 5**.

**Figure 6.**
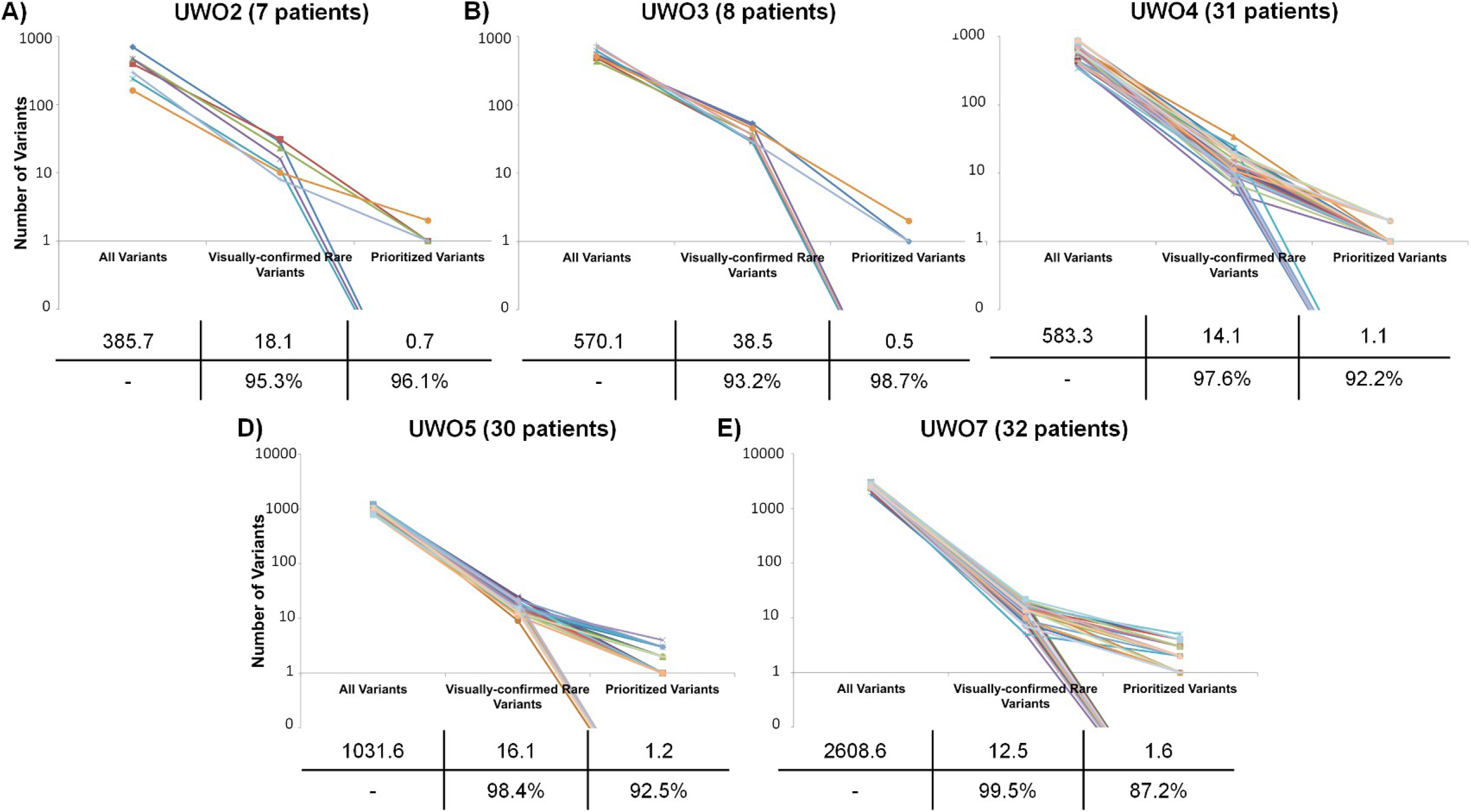
Ladder Plot Representing Variant Identification and Prioritization. Each line is representative of a different sample in each sequencing run (A-E), illustrating the number of unique variants at important steps throughout the variant prioritization process. The left-most point indicates the total number of unique variants. The second point represents the number of unique variants remaining after common (> 5 patients within cohort and/or ≥ 1.0% allele frequency) and false-positive variants were removed. The right-most point represents the final number of unique. No variants were prioritized in the following patients: 2–1A, 2–5A, 2–6A, 3–2A, 3–3A, 3–4A, 3–5A, 3–8A, 4–1B, 4–2C, 4–2F, 4–3B, 4–3D, 4–4B, 4–4E, 5–1G, 5–1H, 5–3D, 5- 4C, 5–4D, 5–4F, 5–4G, 5–4H, 7–1B, 7–1C, 7–1D, 7–1H, 7–2B, 7–2C, 7–2H, 7–3H, 7–4A, 7–4D, 7–4H. The average number of variants per patient at each step is indicated in a table below each plot, along with the percent reduction in variants from one step to another.

**Table 5.** Summary of Prioritized Variants by Gene.

|  | Indel | Nonsense | Missense | Natural Splicing | Cryptic Splicing | Pseudoexon | SR Factor | TF | UTR Structure | UTR Binding | Total |
| --- | --- | --- | --- | --- | --- | --- | --- | --- | --- | --- | --- |
| *ATM* | 0 | 0 | 14 | 2 | 0 | 0 | 18 | 0 | 0 | 1 | 34<sup>‡</sup> |
| *BRCA1* | 2 | 0 | 2 | 0 | 0 | 1 | 7 | 1 | 0 | 0 | 13 |
| *BRCA2* | 0 | 2 | 3 | 0 | 0 | 2 | 4 | 0 | 0 | 0 | 11 |
| *CDH1* | 0 | 0 | 2 | 0 | 0 | 2 | 1 | 1 | 1 | 1 | 8 |
| *CHEK2* | 0 | 0 | 2 | 1 | 0 | 0 | 3 | 0 | 0 | 2 | 6<sup>‡</sup> |
| *PALB2* | 1 | 2 | 3 | 0 | 0 | 0 | 3 | 1 | 0 | 0 | 10 |
| *TP53* | 0 | 0 | 1 | 0 | 0 | 0 | 0 | 2 | 0 | 2 | 5 |

<sup>‡</sup>Counts represent the number of unique variants identified (i.e. a variant is not counted twice if it appeared in multiple individuals).
Three variants were prioritized under multiple categories: *ATM* chr11:108121730A>G (missense and SRFBS), *CHEK2* chr22:29121242G>A (missense, UTR binding), and *CHEK2* chr22:29130520C>T (missense, UTR binding).

Three prioritized variants have multiple predicted roles: *ATM* c.1538A>G in missense and SRFBS, *CHEK2* c.190G>A in missense and UTR binding, and *CHEK2* c.433C>T in missense and UTR binding. Of the 102 patients that we sequenced, 72 (70.6%) exhibited at least one prioritized variant, and some patients harbored more than one prioritized variant (N=33; 32%). **Table S11 (Additional file 14)** presents a summary of all flagged and prioritized variants for patients with at least one prioritized variant.

### Variant Verification

We verified prioritized protein-truncating (N=7) and splicing (N=4) variants by Sanger sequencing (**Table 4** and **Table 2**, respectively). In addition, two missense variants (*BRCA2* c.7958T>C and *CHEK2* c.433C>T) were re-sequenced, since they are indicated as likely pathogenic/pathogenic in ClinVar (**Table S9**). All protein-truncating variants were confirmed, with one exception (*BRCA2* c.7558C>T, no evidence for the variant was present for either strand). Two of the mRNA splicing mutations were confirmed on both strands, while the other two were confirmed on a single strand (*ATM* c.6347+1G>T and *ATM* c.1066–6T>G). Both documented pathogenic missense variants were also confirmed.

## DISCUSSION

NGS technology offers advantages in throughput and variant detection [116], but the task of interpreting the sheer volume of variants in complete gene or genome data can be daunting. The whole genome of a Yoruban male contained approximately 4.2 million SNVs and 0.4 million structural variants [150]. The variant density in the present study (average 948 variants per patient) was 5.3-fold lower than the same regions in HapMap sample NA12878 in Illumina Platinum Genomes Project (5,029 variants) [151]. The difference can be attributed primarily to the exclusion of polymorphisms in highly repetitive regions in our study.

Conventional coding sequence analysis, combined with an IT-based approach for regulatory and splicing-related variants, reduced the set to a manageable number of prioritized variants. Unification of non-coding analysis of diverse protein-nucleic acid interactions using the IT framework accomplishes this by applying thermodynamic-based thresholds to binding affinity changes and by selecting the most significant binding site information changes, regardless of whether the motifs of different factors overlap.

Previously, rule-based systems have been proposed for variant severity classification [152, 153]. Functional validation and risk analyses of these variants are a prerequisite to classification, but this would not be practical to accomplish without first limiting the subset of variants analyzed. With the exception of some (but not all [83]) protein truncating variants, classification is generally not achievable by sequence analysis alone. Only a minority of variants with extreme likelihoods of pathogenic or benign phenotypes are clearly delineated because only these types of variants are considered actionable [152, 153]. The proposed classification systems preferably require functional, co-segregation, and risk analyses to stratify patients. Nevertheless, the majority of variants are VUS, especially in the case of variants occurring beyond exon boundaries. Of the 5,713 variants listed in the BIC database, the clinical significance of 4,102 BRCA1 and BRCA2 variants are either unknown (1,904) or pending (2,198), while 1,535 are classified as pathogenic (Class 5) [154]. Our results cannot be considered equivalent to validation, which might include expression assays [31] or the use of RNA-seq data [155] (splicing), qRT-PCR [156] (transcription), SHAPE analysis (mRNA 2° structure) [35], and binding assays to determine functional effects of variants. Other post-transcriptional processes (eg. miRNA regulation) affected by variants have not been addressed in this study, but should also be amenable to IT-based modeling. With the proposed approach, functional prediction of variants could precede or at least inform the classification of VUS.

It is unrealistic to expect all variants to be functionally analyzed, just as it may not be feasible to assess family members for a suspected pathogenic variant detected in a proband. The prioritization procedure reduces the chance that significant variants have been overlooked. Capturing coding and non-coding regions of HBOC-related genes, combined with the framework for assessing variants balances the need to comprehensively detect all variation in a gene panel with the goal of identifying variants likely to be phenotypically relevant.

### Non-coding variants

Variant density in non-coding regions significantly exceeded exonic variants by > 60-fold, which, in absolute terms, constituted 1.6% of the 15,311 variants. This is comparable to whole genome sequencing studies, which typically result in 3–4 million variants per individual, with < 2% occurring in protein coding regions [157]. IT analysis prioritized 3 natural SS, 36 SRFBS, 5 TFBS, and 6 RBBS variants and 5 predicted to create pseudoexons. Two SS variants in *ATM* (c.3747–1G>A and c.6347+1G>T) were predicted to completely abolish the natural site and cause exon skipping. A *CHEK2* variant (c.320–5A>T) was predicted to result in leaky splicing.

The IT-based framework evaluates all variants on a common scale, based on bit values, the universal unit that predicts changes in binding affinity [158]. A variant can alter the strength of one or a “set” of binding sites; the magnitude and direction of these changes is used to rank their significance. The models used to derive information weight matrices take into account the frequency of all observed bases at a given position of a binding motif, making them more accurate than consensus sequence and conservation-based approaches [31].

IT has been widely used to analyze natural and cryptic SSs [31], but its use in SRFBS analysis was only introduced recently [80]. For this reason, we assigned conservative, minimum thresholds for reporting information changes. Although there are examples of disease-causing variants resulting in small changes in *R_i_* [159–166], the majority of deleterious splicing mutations that have been verified functionally, produce large information changes. Among 698 experimentally deleterious variants in 117 studies, only 1.96% resulted in < 1.0 bit change [31]. For SRFBS variants, the absolute information changes for deleterious variants ranged from 0.2 - 17.1 bits (mean 4.7 ± 3.8). This first application of IT in TFBS and RBBS analysis, however, lacks a large reference set of validated mutations for the distribution of information changes associated with deleterious variants. The release of new ChIP-seq datasets will enable IT models to be derived for TFs currently unmodeled and to improve existing models [167].

Pseudoexon activation results in disease-causing mutations [168], however such consequences are not customarily screened for in mRNA splicing analysis. IT analysis was used to detect variants that predict pseudoexon formation and 5 variants were prioritized. Previously, we have predicted experimentally proven pseudoexons with IT (Ref 34: Table 2, No #2 and Ref 169: Table 2, No #7) [34, 169]. Although it was not possible to confirm prioritized variants in the current study predicted to activate pseudoexons because of their low allele frequencies, common intronic variants that were predicted to form pseudoexons were analyzed. We then searched for evidence of pseudoexon activation in mapped human EST and mRNA tracks [170] and RNA-seq data of breast normal and tumour tissue from the Cancer Genome Atlas project [171]. One of these variants (rs6005843) appeared to splice the human EST HY160109 [172] at the predicted cryptic splice site and is expressed within the pseudoexon boundaries.

Variants that were common within our population sample (i.e. occurring in > 5 individuals) and/or common in the general population (> 1.0% allele frequency) reduced the list of flagged variants substantially. This is now a commonly accepted approach for reducing candidate disease variants [152], based on the principle that the disease-causing variants occur at lower population frequencies. Variants occurring in > 5 patients all either had allele frequencies above 1.0% or, as shown previously, resulted in very small Δ*R_i_,* values [173].

The genomic context of sequence changes can influence the interpretation of a particular variant [31]. For example, variants causing significant information changes may be interpreted as inconsequential if they are functionally redundant or enhancing existing binding site function (see *IT-Based Variant Analysis* for details). Our understanding of the roles and context of these cognate protein factors is incomplete, which affects confidence in interpretation of variants that alter binding. Also, certain factors with important roles in the regulation of these genes, but that do not bind DNA directly or in a sequence-specific manner, (eg. CtBP2 [174]), could not be included. Therefore, some variants may have been incorrectly excluded.

### Coding sequence changes

We also identified 4 nonsense and 3 indels in this cohort. In one individual, a 19 nt *BRCA1* deletion in exon 15 causes a frameshift leading to a stop codon within 14 codons downstream. This variant, rs80359876, is considered clinically relevant. Interestingly, this deletion overlaps two other published deletions in this exon (rs397509209 and rs80359884). This raises the question as to whether this region of the *BRCA1* gene is a hotspot for replication errors. DNA folding analysis indicates a possible 15 nt long stem-loop spanning this interval as the most stable predicted structure (data not shown). This 15 nt structure occurs entirely within the rs80359876 and rs397509209 deletions and partially overlaps rs80359884 (13 of 15 nt of the stem loop). It is plausible that the 2° structure of this sequence predisposes to a replication error that leads to the observed deletion.

Missense coding variants were also assessed using multiple *in silico* tools and evaluated based on allele frequency, literature references, and gene-specific databases. Of the 27 prioritized missense variants, the previously reported *CHEK2* variant c.433G>A (chr22:29121242G>A; rs137853007) stood out, as it was identified in one patient (4–3C.5–4G) and is predicted by all 4 *in silico* tools to have a damaging effect on protein function. Accordingly, Wu et al. (2001) demonstrated reduced *in vitro* kinase activity and phosphorylation by ATM kinase compared to the wild-type protein [175], presumably due to the variant’s occurrence within the forkhead homology-associated domain, involved in protein-phosphoprotein interactions [176]. Implicated in Li-Fraumeni syndrome, known to increase the risk of developing several types of cancer including breast [177, 178], this variant is expected to result in a misfolded protein that would be targeted for degradation via the ubiquitin-proteosome pathway [179]. Another important missense variant is c.7958T>C (chr13:32936812T>C; rs80359022; 4–4C) in exon 17 of *BRCA2.* Although classified as being of unknown clinical importance in both BIC and ClinVar, it has been classified as pathogenic based on posterior probability calculations [180].

It is unlikely that all prioritized variants are pathogenic in patients carrying more than one prioritized variant. Nevertheless, a polygenic model for breast cancer susceptibility, whereby multiple moderate and low-risk alleles contribute to increased risk of HBOC may also account for multiple prioritized variants [181, 182]. There was a significant fraction of patients (29.4%) in whom no variants were prioritized. This could be due to: a) the inability of the analysis to predict a variant affecting the binding sites analyzed, b) a pathogenic variant affects a function that was not analyzed or in a gene that was not sequenced, or c) the significant family history was not due to heritable, but instead to shared environmental influences.

*BRCA* coding variants were found in individuals who were previously screened for lesions in these genes, suggesting this NGS protocol is a more sensitive approach for detecting coding changes. However, the previous testing was predominantly based on PTT and MLPA methods, which have lower sensitivity than sequence analysis. Nevertheless, we identified 2 *BRCA1* and 2 *BRCA2* variants predicted to encode prematurely truncated proteins. Fewer non-coding *BRCA* variants were prioritized (15.7%) than expected by linkage analysis [37], however this presumes at least 4 affected breast cancer diagnoses per pedigree, and, in the present study, the number of affected individuals per family was not known.

Prioritization of a variant does not equate with pathogenicity. Some prioritized variants may not increase risk, but may simply modify a primary unrecognized pathogenic mutation. A patient with a known *BRCA1* nonsense variant, used as a positive control, was also found to possess an additional prioritized variant in *BRCA2* (missense variant chr13:32911710A>G), which was flagged by PROVEAN and SIFT as damaging, as well as flagged for changing an SRFBS for abolishing a PTB site (while simultaneously abolishing an exonic hnRNPA1 site). This variant has been identified in cases of early onset prostate cancer and is considered a VUS in ClinVar [133]. A larger cohort of patients with known pathogenic mutations would be necessary to calculate a background/basal rate of falsely flagged variants.

Other groups have attempted to develop comprehensive approaches for variant analysis, analogous to the one proposed here [183–185]. While most employ high-throughput sequencing and classify variants, either the sequences analyzed or the types of variants assessed tend to be limited. In particular, non-coding sequences have not been sequenced or studied to the same extent, and none of these analytical approaches have adopted a common framework for mutation analysis.

Our published oligonucleotide design method [64] produced an average sequence coverage of 98.8%. The capture reagent did not overlap conserved highly repetitive regions, but included divergent repetitive sequences. Nevertheless, neighboring probes generated reads with partial overlap of repetitive intervals. As previously reported [135], we noted that false positive variant calls within intronic and intergenic regions were the most common consequence of dephasing in low complexity, pyrimidine-enriched intervals. This was not alleviated by processing data with software programs based on different alignment or calling algorithms. Manual review of all intronic or intergenic variants became imperative. As these sequences can still affect functional binding elements detectable by IT analysis (i.e. 3’ SSs and SRFBSs), it may prove essential to adopt or develop alignment software that explicitly and correctly identifies variants in these regions [135]. Most variants were confirmed with Sanger sequencing (10/13), and those that could not be confirmed are not necessarily false positives. A recent study demonstrated that NGS can identify variants that Sanger sequencing cannot, and reproducing sequencing results by NGS may be worthwhile before eliminating such variants [186].

## CONCLUSIONS

Through a comprehensive protocol based on high-throughput, IT-based and complementary coding sequence analyses, the numbers of VUS can be reduced to a manageable quantity of variants, prioritized by predicted function. Exonic variants corresponded to a small fraction of prioritized variants, illustrating the importance of sequencing non-coding regions of genes. We propose that our approach for variant flagging and prioritization is an intermediate bridge between high-throughput sequencing, variant detection, and the time-consuming process of variant classification, including pedigree analysis and functional validation.

## AVAILABILITY OF SUPPORTING DATA

Variants will be deposited with the ENIGMA Consortium (www.enigmaconsortium.org), which is a designated organization for curation of HBOC mutations and which is charged with protection of genetic privacy of participants.

## LIST OF ABBREVIATIONS

ASSEDA: Automated Splice Site and Exon Definition Analysis
BIC: Breast Cancer Information Core Database
CASAVA: Consensus Assessment of Sequencing and Variation
CIS-BP-RNA: Catalog of Inferred Sequence Binding Preferences of RNA binding proteins
CRAC: Complex Reads Analysis and Classification
DM^2^: Domain Mapping of Disease Mutations
ENIGMA: Evidence-based Network for the Interpretation of Germline Mutant Alleles
ExPASy: Expert Protein Analysis System
GATK: Genome Analysis Toolkit
HBOC: Hereditary Breast and Ovarian Cancer
HGMD: Human Gene Mutation Database
IARC: International Agency for Research on Cancer
IGV: Integrative Genomics Viewer
Indel: Insertion/deletion
IT: Information theory
LOVD: Leiden Open Variant Database
MGL: Molecular Genetics Laboratory
MLPA: Multiplex Ligation Probe Amplification
NGS: Next-Generation Sequencing
PTB: Polypyrimidine tract binding protein
PTT: Protein Truncation Test
PWM: Position Weight Matrix
RBBS: RNA-Binding protein Binding Site
RBP: RNA-Binding Protein
RBPDB: RNA-Binding Protein DataBase
*R_i_*: Individual information
*R_sequence_*: Mean information content
SHAPE: Selective 2’-Hydroxyl Acylation analyzed by Primer Extension
SNV: Single Nucleotide Variant
SRF: Splicing Regulatory Factor
SRFBS: Splicing Regulatory Factor Binding Site
SS: Splice Site
TF: Transcription Factor
TFBS: Transcription Factor Binding Site
UTR: Untranslated Region
VCF: Variant Call File
VUS: Variants of Uncertain Significance
Δ*R_i_*: Change in individual information.
Patient Sample IDs are assigned in following manner: number-number+letter (i.e. 1–1A). If a sample was repeated, the IDs are separated by a “.” (i.e. 1–1A.2–1A)

## AUTHORS’ CONTRIBUTION

PKR designed, coordinated, and supervised the study, which was motivated by discussions with JHMK regarding prioritization of VUS. EJM performed probe design and synthesis. EJM and NGC performed sample preparation and sequencing. EJM wrote software and performed bioinformatic analysis. EJM, NGC, and AMP conducted variant analysis and prioritization. RL generated the TFBS information models and EJM generated the RBBS, SRF, and splicing information models. AMP confirmed prioritized variants by Sanger sequencing. MH and AL conducted the SHAPE analysis. EJM, NGC, AMP, JHMK, and PKR wrote the manuscript, which has been approved by all authors.

### Additional files

**Additional file 1: Supplementary Methods**

Format: PDF

**Additional file 2: Provincial Eligibility Criteria.** Risk Categories for Individuals Eligible for Screening for a Genetic Susceptibility to Breast or Ovarian Cancer as determined by the Ontario Ministry of Health and Long Tern-Care Referral Criteria for Genetic Counseling

Format: PDF

**Additional File 3: Table S1.** TFs For Which Information Weight Matrices Were Built And Factor’s Role in Transcription

Format: XLSX

**Additional file 4: Table S2**. UTR Sequences Used for SHAPE Analysis on SNPfold-flagged Variants

Format: XLSX

**Additional file 5: Table S3**. Primer Sequences for Sanger Sequencing of Likely Pathogenic Variants

Format: XLSX

**Additional file 6: Figure S1**. *BRCA1* Deletion Inaccurately Aligned by CASAVA

Format: PDF

**Additional file 7: Table S4**. Variants Identified Within Natural Sites

Format: XLSX

**Additional file 8: Table S5**. Variants Predicted by IT to Affect SRFBSs

Format: XLSX

**Additional file 9: Table S6**. Variants Predicted by IT to Affect TFBSs

Format: XLSX

**Additional file 10: Table S7**. Top Changes in RBBSs Predicted by IT for Variants Predicted to Significantly Alter RNA Structure

Format: XLSX

**Additional file 11: Table S8**. Missense Variants Identified In 6 Patients Or More

Format: XLSX

**Additional file 12: Table S9**. Missense Variants and Their Classification

Format: XLSX

**Additional file 13: Table S1**0. Prioritized Variants by Gene

Format: XLSX

**Additional file 14: Table S1**1.

All Flagged and Prioritized Variants by Patient

Format: XLSX

## ACKNOWLEDGEMENTS

We thank Dr. Peter Ainsworth and Alan Stuart of the MGL, Molecular Diagnostics Division, Dept of Pathology and Laboratory Medicine at the London Health Sciences Centre for access to patient DNA samples and the use of the BioMek^®^ FXP Automation Workstation. Information models for protein binding were computed with PoWeMaGen (Nathan Bryans) and variants were scanned with Mutation Analyzer, a modification of the Shannon Human Splicing Mutation Pipeline (Coby Viner). Edwin Dovigi contributed to design and synthesis of the custom capture reagent. PKR is supported by the Canadian Breast Cancer Foundation, Canadian Foundation for Innovation, Canada Research Chairs Secretariat and the Natural Sciences and Engineering Research Council of Canada (NSERC Discovery Grant 371758–2009). NGC is funded by the CIHR Strategic Training Program in Cancer Research and Technology Transfer (CaRTT) and the Pamela Greenaway-Kohlmeier Translational Breast Cancer Research Unit (TBCRU) awards. Our work was made possible by the facilities of the Shared Hierarchical Academic Research Computing Network (SHARCNET) and Compute/Calcul Canada.

